# Preserved T cell reactivity to the SARS-CoV-2 Omicron variant indicates continued protection in vaccinated individuals

**DOI:** 10.1101/2021.12.30.474453

**Authors:** Lorenzo De Marco, Silvia D’Orso, Marta Pirronello, Alice Verdiani, Andrea Termine, Carlo Fabrizio, Alessia Capone, Andrea Sabatini, Gisella Guerrera, Roberta Placido, Manolo Sambucci, Daniela F. Angelini, Flavia Giannessi, Mario Picozza, Carlo Caltagirone, Antonino Salvia, Elisabetta Volpe, Maria Pia Balice, Angelo Rossini, Olaf Rötzschke, Emiliano Giardina, Luca Battistini, Giovanna Borsellino, Santa Lucia Foundation, Rome, Italy

## Abstract

**Importance:** The emergence of the highly contagious Omicron variant of SARS-CoV-2 and the findings of a significantly reduced neutralizing potency of sera from convalescent or vaccinated individuals imposes the study of cellular immunity to predict the degree of immune protection to the yet again new coronavirus.

**Design:** Prospective monocentric observational study.

**Setting:** Conducted between December 20-21 at the Santa Lucia Foundation IRCCS.

**Participants:** 61 volunteers (Mean age 41.62, range 21-62; 38F/23M) with different vaccination and SARS-CoV-2 infection backgrounds donated 15 ml of blood. Of these donors, one had recently completed chemotherapy, and one was undergoing treatment with monoclonal antibodies; the others reported no known health issue.

**Main Outcome(s) and Measure(s):** The outcomes were the measurement of T cell reactivity to the mutated regions of the Spike protein of the Omicron SARS-CoV-2 variant and the assessment of remaining T cell immunity to the spike protein by stimulation with peptide libraries.

**Results:** Lymphocytes from freshly drawn blood samples were isolated and immediately tested for reactivity to the Spike protein of SARS-CoV-2. T cell responses to peptides covering the mutated regions in the Omicron variant were decreased by over 47% compared to the same regions of the ancestral vaccine strain. However, overall reactivity to the peptide library of the full-length protein was largely maintained (estimated 83%). No significant differences in loss of immune recognition were identified between groups of donors with different vaccination and/or infection histories.

**Conclusions and Relevance:** We conclude that despite the mutations in the Spike protein, the SARS-CoV-2 Omicron variant is nonetheless recognized by the cellular component of the immune system. It is reasonable to assume that protection from hospitalization and severe disease is maintained.

**Key Points:** *Question:* Does the Omicron variant of SARS-CoV-2 escape cellular immunity?

*Findings:* This observational study was performed on 61 vaccinated donors with established immunity to SARS-CoV-2. Cellular responses to the mutated regions of the Omicron Spike protein were detected in 80% of donors. The mutations reduced T cell recognition by 47% compared to the vaccine strain. Reactivity to the whole Spike protein, however, was present in 100% of donors, and the fraction of remaining immunity to SARS-CoV-2 was estimated to be 83%.

*Meaning:* Cellular immunity to the Omicron variant is maintained despite the mutations in its Spike protein, and may thus confer protection from severe COVID-19 in vaccinated individuals.

## Introduction

Far from being weakened, the pandemic of severe acute respiratory syndrome coronavirus 2 (SARS-CoV-2) has found new strength in another wave of infections with the Omicron variant ^1^. New variants represent a threat, since their mutations may confer both increased transmissibility and ability to evade previously established immune responses. Mutations in the RBD region correlate with the lower neutralization potency of sera collected from convalescent or vaccinated individuals ^2^, setting the stage for immune evasion by the mutated virus. Conveniently, T cell responses are characterized by vast cross-reactivity ^3^, and cellular immunity is maintained also in face of mutations which determine the escape from antibody recognition ^4^. Still, infection of a fully vaccinated individual with a slightly different version of the immunizing pathogen finds a situation of “pre-existing” immunity, mostly mediated by the cellular component of the immune response. This has shown to be the case in the setting of another respiratory virus vaccine, the adjuvanted clade 1 H5N1 subunit vaccine for avian influenza ^5^, where boosting determined the expansion of a memory H5-specific T cell population that could also recognize H5 from other clades. Will the intrinsic cross-reactivity of the Spike-specific T cells induced by vaccination and/or infection confer a broad enough repertoire for them to quickly respond to SARS-CoV-2 infection with emerging variants? Studies on the fine specificity and on the persistence of Spike-specific T cells in COVID-19 convalescents have indicated that the cellular response to SARS-CoV-2, far from being monoclonal, is composed by clones recognizing multiple sites of the RBD region, thus offering broad reactivity against Spike epitopes ^6,7^, and induction of broadly reactive CD4^+^ and CD8^+^ memory cells has been shown to occur following infection and/or vaccination ^8^. Epidemiological data shows striking increases in SARS-CoV-2 infection rates and a very rapid spread worldwide of the Omicron variant, emerged in November 2021 ^9^. Here, we investigate the T cell response to the mutated regions of the Spike protein from the Omicron variant in 61 vaccinated individuals. By also measuring reactivity to the equivalent regions from the ancestral vaccine strain and to the whole Spike protein, we estimate the degree of remaining immunity to the SARS-CoV-2 spike protein.

## Methods

### Study Design

This observational study was carried out on December 20-21 at the Santa Lucia Foundation Hospital and was approved by the local Ethics Committee. After providing informed consent, 61 volunteers selected among hospital workers, scientists, and their families and acquaintances (Mean age 41.62, range 21-62; 38F/23M) donated 15 ml of blood. Donors were divided in 5 groups, based on their vaccination/infection history: 1) 2D: 2 doses of vaccine; 2) 3D: 3 doses of mRNA vaccine; 3) Het: heterologous vaccination with adenoviral vector followed by mRNA vaccine; 4) VAC-COV: vaccinated individuals who had subsequently been infected (all asymptomatic or with mild disease); 5) COV-VAC: COVID-19-recovered individuals who had subsequently been vaccinated (Table 1).

**Tab.1.** Donor characteristics. *= Infected with SARS-CoV2 Omicron variant. P = BNT162b2 Pfizer/BioNtech vaccine. M = mRNA-1273 Moderna vaccine. J = Ad26.COV2.S Janssen vaccine. AZ = ChAdOx1-S AstraZeneca vaccine

| Immunological History | Subjects | Age<br>Mean<br>(range) | Sex<br>M/F | Vaccine<br>(in chronological order) |
| --- | --- | --- | --- | --- |
| 2D<br>(10) | 6 | 42.5<br>(23 - 56) | 3/7 | 2 P |
|  | 3 |  |  | 2 M |
|  | 1 |  |  | 2 AZ |
| 3D<br>mRNA<br>(15) | 14 | 52<br>(26 - 60) | 5/10 | 3 P |
|  | 1 |  |  | 2 P + 1 M |
| Heterologous<br>(11) | 3 | 53<br>(26 - 62) | 3/8 | 2 AZ + 1 M |
|  | 6 |  |  | 2 AZ + 1 P |
|  | 2 |  |  | 1 J + 1 P |
| COVID-19<br>then<br>Vaccine<br>(12) | 7 | 53.5<br>(22 - 60) | 8/4 | 2 P |
|  | 2 |  |  | 3 P |
|  | 1 |  |  | 1 J |
|  | 1 |  |  | 1 M |
|  | 1 |  |  | 1 J + 1 M |
| Vaccine<br>then<br>COVID-19<br>(13) | 9 | 37<br>(21 - 54) | 4/9 | 2 P |
|  | 1 |  |  | 1 J |
|  | 1 |  |  | 2 AZ + 1 M |
|  | 1* |  |  | 2 AZ |
|  | 1* |  |  | 2 P |

Peripheral blood mononuclear cells (PBMCs) were isolated and immediately tested in an in vitro assay consisting of incubation with 3 different peptide pools with a peptide length of 15 amino acids: 1) an overlapping peptide pool spanning the entire Spike protein from the ancestral vaccine strain (PoolS); 2) a peptide pool covering only the mutated regions of the Spike protein from the Omicron variant (Pool_Mut_); 3) a peptide pool covering the same regions as above, but from the ancestral strain (Pool_Ref_) (all pools 1 μg/ml each, Miltenyi Biotec). After 18 hours, cells were stained with fluorochrome-conjugated monoclonal antibodies for the detection of the expression of surface Activation Induced Markers (AIM) in CD4+ and CD8+ T cell subsets (Table 2), and supernatants were collected for measurement of IFN-γ release by ELISA (R&D Systems). Activated CD4+ cells were defined as CD40L+CD69+, while the CD137+CD69+ fraction identified activated CD8+ cells, as previously described ^10^. After staining, cells were acquired on an Aurora (Cytek) or on a Cytoflex LS (Beckman Coulter) flow cytometers. In parallel, 50 μl of corresponding whole blood samples were stained with anti CD3, anti CD4, and anti CD8 for the determination of absolute cell counts. Data were analyzed with FlowJo v 10.8. Statistics were performed as detailed in Suppl. Methods, and data were visualized using GraphPad PRISM v.9.

**Tab.2.** Antibodies used.

| Antibody | Fluorochrome | Clone | Company | Titer |
| --- | --- | --- | --- | --- |
| CD14 | FITC | REA599 | Miltenyi | 1:100 |
| CD19 | FITC | MAB-1118F | ImmunolScience | 1:30 |
| CD69 | BB700 | FN50 | Bect.Dick. | 1:100 |
| CD154 | PE | 24-31 | eBioscience | 1:30 |
| CD4 | PECy5.5 | 13B8.2 | Coulter | 1:100 |
| CD8 | BV605 | SK1 | Bect. Dick. | 1:40 |
| CD137 | BUV395 | 4B4-1 | Bect. Dick. | 1:30 |
| CD3 | BUV496 | UCHT1 | Bect.Dick. | 1:50 |
| LD | Near-IR |  | ThermoFisher | 1:60 |

## Results

First, we established T cell responsiveness to the Spike protein by utilizing a FACS-based activation induced marker (AIM) assays to quantify SARS-CoV-2-specific CD4+ and CD8+ T cells in a cohort of 61 vaccinated donors (Fig.1A). The frequency of donors showing CD4+ and CD8+ T cell reactivity to the overlapping peptide pool covering the whole Spike protein (PoolS) was 100% and 96%, respectively (Fig. 1B). CD4+ and CD8+ T cell reactivity to the selected peptide pools covering the mutated regions of the omicron variant (Pool_Mut_) was detected in 74% and 60% of donors, respectively. Recognition of the corresponding unmutated regions (Pool_Ref_) by CD4+ and CD8+ cells was present in in 96% and 63% of donors, respectively. Within the different groups of donors, a larger fraction of individuals who had received the heterologous regimen of vaccination showed T cell reactivity (91% for CD4+ and 73% for CD8+) to the Pool_Mut_. On the other hand, fewer individuals who had received 2 doses of vaccine responded to the Pool_Mut_ (70% for CD4+ and 30% for CD8). However, probably also due to the small sample size, these differences were not statistically significant.

**Fig. 1.**
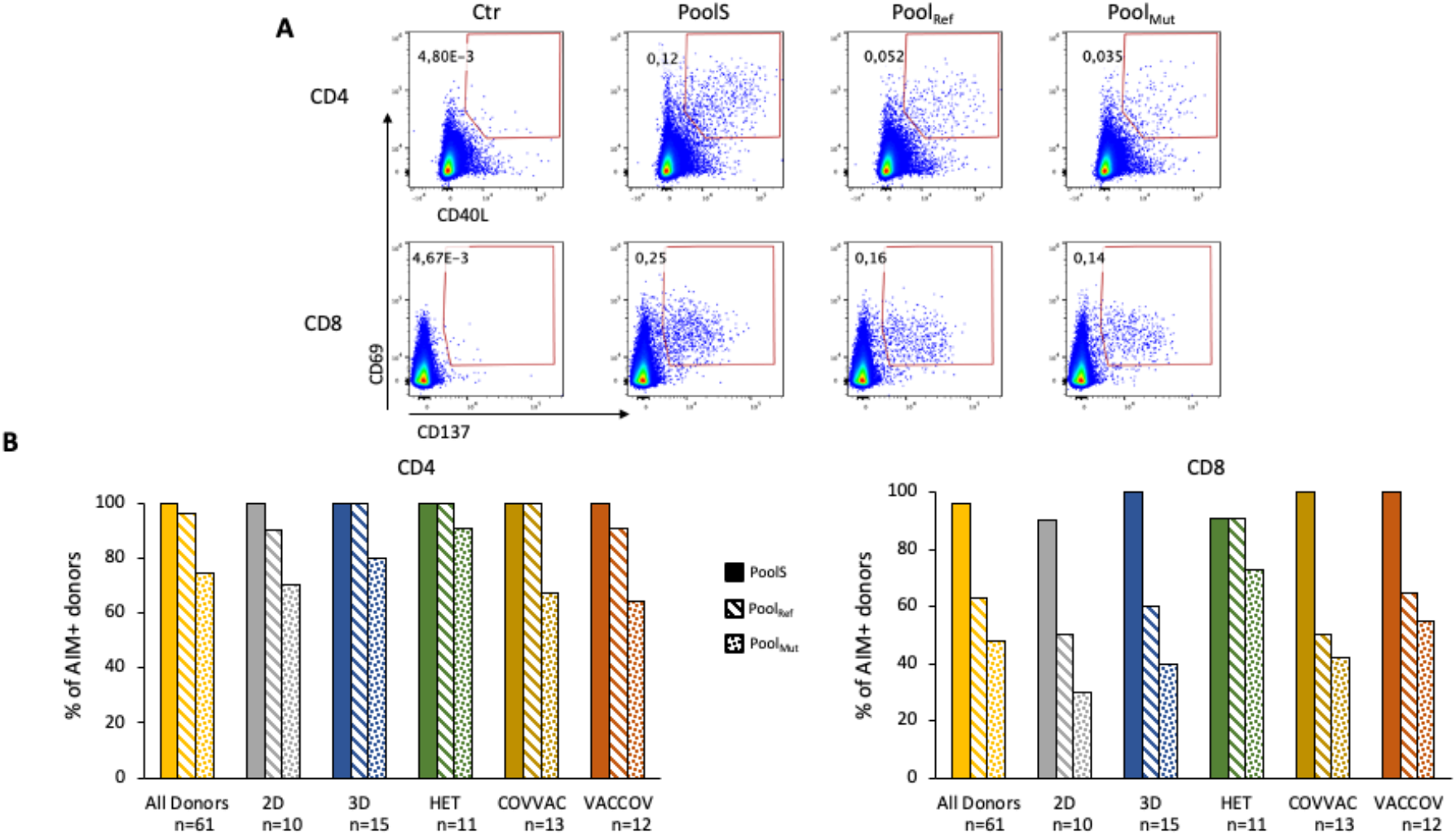
T cell responses to the Spike protein. A) Representative flow cytometry plots gated on CD4+ or CD8+ T cells showing up-regulation of activation markers (CD69 and CD40L for CD4+ cells and CD69 and CD137 for CD8 cells) following overnight stimulation with a pool of overlapping peptides covering the whole Spike protein from the ancestral vaccine strain (Pools), a peptide pool covering only the mutated regions of the Spike protein from the Omicron variant (PoolMut) or a peptide pool covering the same regions as above, but from the ancestral strain (PoolRef). B) percentage of individuals in each group presenting CD4+ (left) and CD8+ (right) Spike-specific responses.

Overall, the fraction of CD4+ and CD8+ T cells activated by Pool_Mut_ was significantly lower compared to that of T cells recognizing the same regions from the Wuhan strain used for vaccine design (median 0,109 % PoolS and 0,039% Pool_Mut_ for CD4+ T cells, p = 0,0001, and 0,039% PoolS, 0,02% Pool_Mut_ for CD8 cells, p = 0,0001) (Fig 2A and B). The reduction in T cell numbers reactive to the omicron variant was found to be significant in all groups of donors, regardless of vaccination and SARS-CoV-2 infection history. Although not significant, the group of donors who had received heterologous vaccination showed a lower reduction in Spike-protein recognition both in the CD4+ and in the CD8+ T cell subset. Individuals who had been infected despite vaccination also showed a lower, albeit not significant, reduction in the fraction of CD8+ T cells recognizing the mutated regions of Spike. CD8+ T cell reactivity, when present, was more conserved compared to the CD4+ subset.

**Fig. 2.**
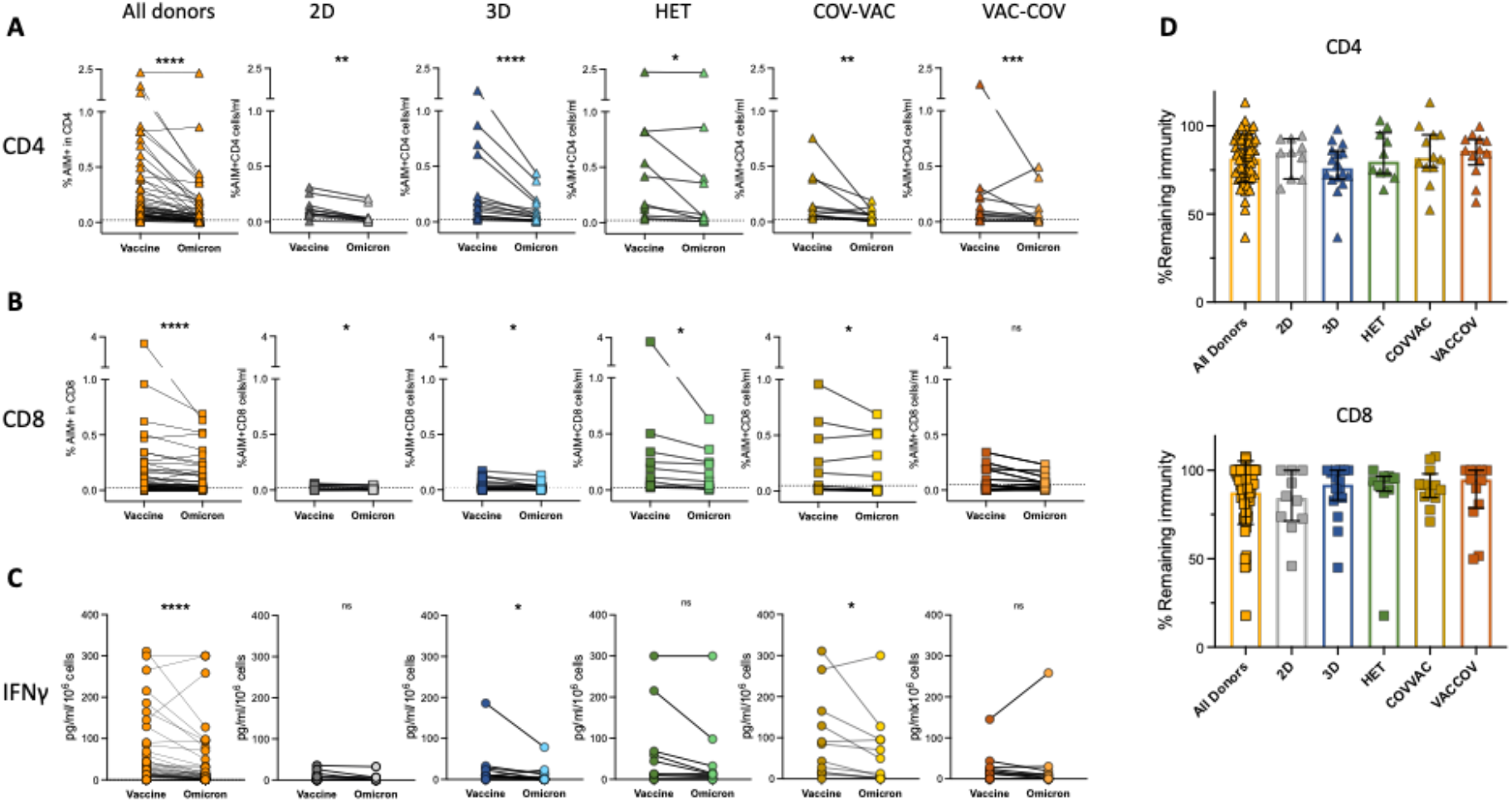
Reduced recognition of mutated regions of the Spike protein in the Omicron variant. Freshly isolated lymphocytes were incubated with peptide pools encompassing the mutated regions of the Spike protein in the Omicron variant (Omicron), and with the reference peptide pool of the same region in the ancestral vaccine strain (Vaccine). Activated CD4+ (A) (CD69+CD40L+) and CD8+(B) (CD69CD137+) cells were identified by flow cytometry. Background T cell activation in paired unstimulated cultures was subtracted. Dotted lines indicate the threshold for positivity (median-75th percentile of values obtained in unstimulated cultures). 2D: = 2 doses of vaccine: 3D: 3 doses mRNA vaccine, HET: 1-2 doses adenoviral vector vaccine + 1 dose mRNA, COV-VAC: covid convalescents subsequently vaccinated; VAC-COV Vaccinated individuals subsequently infected with SARS-CoV-2. Differences were assessed using Friedman rank sum test with Omicron exposure (vaccine/omicron) and subject ID as foved and random effects respectively. P<0.05, **P<0.01***P<0.001; ****P<0.0001; N.S. Not Significant C) IFN-y production measured in the suprnatants of the peptide-stimulated cultures. D) Fraction of preserved CD4+ (top) and CD8+ (bottom) reactivity to the whole Spike protein in each group, after having substracted loss of reactivity to the mutated regions.

IFN-γ released in the cultures was also significantly reduced when cells were incubated with the peptide pools covering the mutated regions of Spike (p = 0,001), compared to pools spanning the equivalent regions of the ancestral strain (Fig.2C). Thus, this data shows that CD4+ and CD8+ T cell activation and function are affected by the mutations of the Spike protein carried by the Omicron variant.

While we measure a significant reduction in T cell reactivity to the mutated regions of the Spike protein from the Omicron variant, these account for a small fraction of the total protein. To estimate the remaining immunity we subtracted the difference in the T cell response against the mutated and unmutated Pools from the response against the complete protein pool ( (AIM_PoolS_ - (AIM_PoolRef_ - AIM_PoolMut_)). With this simple calculation we estimate that overall T cell reactivity to the Spike protein of the Omicron variant is maintained by 83% (Fig. 2D), with no significant differences between the different groups of donors. Thus, this data shows that regardless of vaccination and/or infection history, T cell responses to the Omicron variant are maintained.

## Discussion

Compared to the Wuhan strain, the Omicron variant carries over 35 mutations in the Spike protein. The impact of these mutations on antibody recognition has been shown to be substantial, with a significant loss of neutralizing activity in serum from both convalescent and vaccinated individuals ^2,11^. In previous variants, although antibody neutralizing potency was decreased, T cell responses were maintained ^4,12^, although some mutations have been shown to affect CD8 T cell recognition ^13^. Here, we find that T cell responses against the mutated regions in Omicron are strongly affected. However, as these regions cover only a small fraction of the whole protein the overall response against Omicron Spike appears to be largely preserved. Reactivity to the variant was similar regardless of vaccination and/or infection history. The finding of persisting and robust T cell responses despite the mutations in Omicron thus provide confidence that the cellular immunity against this variant will not be compromised.

## Acknowledgments

We thank the volunteers for donating their blood and time, and the nurses for their assistance. We thank Enrico Ghersi and the Cytek team for their support, and Mirko Lanuti and Claudia Maldini from Miltenyi for their collaboration.

