## Supplementary Methods for "Preserved T cell reactivity to the SARS-CoV-2 Omicron variant indicates continued protection in vaccinated individuals"

Biostatistical Analysis. The differences between groups in CD4 and CD8 activated T cells were assessed for each experimental condition (PoolS, $Pool_{Ref}$ and $Pool_{Mut}$) using multiple Kruskal-Wallis rank sum tests. Pairwise post-hoc comparisons were performed using the Wilcoxon rank sum test with False Discovery Rate (FDR) correction for multiple testing. Within-groups differences in CD4+ and CD8+ were assessed in $Pool_{Ref}$ and $Pool_{Mut}$ conditions using Friedman rank sum test with omicron exposure and subject ID as fixed and random effects respectively. Kendall's W was used to compute effect size following the Cohen's interpretation guidelines of 0.1 - < 0.3 (small effect), 0.3 - < 0.5 (moderate effect) and >= 0.5 (large effect). The obtained effect-sizes were used in multiple two-tailed post-hoc power analyses for dependent means to estimate the obtained statistical power with $\boldsymbol{\alpha} = 0.05$. The slopes of the regression line between $Pool_{Ref}$ and $Pool_{Mut}$ were computed for $CD4_{AIM}$ and $CD8_{AIM}$ for each subject. The obtained slopes were compared between groups using multiple Kruskal-Wallis rank sum tests and Heteroscedastic one-way ANOVA for medians with FDR correction. All the statistical analyses were performed using the following R(v.4.1.2) libraries: "lme4", "lmerTest", "WRS2".
